# PSMA-specific CAR-engineered macrophages for therapy of prostate cancer

**DOI:** 10.1101/2024.09.07.611792

**Authors:** Yangli Xu, Duoli Xie, Chunhao Cao, Yue Ju, Xinxin Chen, Lili Guan, Xuelong Li, Luo Zhang, Chao Liang, Xiushan Yin

## Abstract

Chimeric antigen receptor (CAR)-modified macrophages (CAR-Ms) are a promising approach for the treatment of solid tumors due to its high infiltration and immune-regulation activity. Prostate cancer is a typical solid tumor associated with highly immunosuppressive microenvironment. To date, the potential application of CAR-M cell therapy in prostate cancer has been infrequently explored. The prostate-specific membrane antigen (PSMA) functions as a specific biomarker for prostate cancer. In this study, we assessed the antitumor efficacy of PSMA-targeted CAR-Ms in preclinical models. CAR-Ms were engineered to express a PSMA-specific single-chain variable fragment (scFv) and co-stimulatory domains. *In vitro* data demonstrated specific cytotoxicity of CAR-Ms against PSMA-expressing prostate cancer cells, which was further supported by transcriptome analysis demonstrating the pro-inflammatory phenotypes of CAR-Ms. *In vivo* studies using xenograft mouse models confirmed significant tumor regression after administration of PSMA-targeted CAR-Ms compared to controls. Histopathological analysis showed infiltration of CAR-Ms into tumor tissues without off-target toxicity. These results highlight the strong antitumor activity and safety of PSMA-targeted CAR-Ms, supporting their potential as a new immunotherapy for prostate cancer.

## Introduction

Prostate cancer is a malignant neoplasm arising from the epithelial tissue of the prostate gland and ranks among the most common cancers in the male genitourinary system^[1, 2]^. The Global Cancer Statistics 2024 report indicates a yearly increase in the incidence of prostate cancer by 2%-3% from 2015 to 2019, positioning it as the second most prevalent cancer in men and responsible for 29% of new cancer cases in this demographic^[3]^. Traditional therapeutic approaches, including surgical resection, radiation therapy, and androgen deprivation therapy, have been the cornerstone of prostate cancer management^[4]^. Despite significant advancements, these conventional approaches carry inherent limitations. Surgical interventions come with the risk of postoperative complications such as incontinence and sexual dysfunction, while radiation and hormone therapies may lead to cumulative toxicities and treatment resistance^[5-8]^. Moreover, these treatments often prove less effective against metastatic prostate cancer, underscoring an urgent need for innovative therapeutic strategies that can overcome these challenges.

The dawn of immunotherapy, which harnesses the body’s own immune system to identify and eliminate cancer cells, has initiated a transformative era in oncological care, offering new avenues of hope where traditional treatments have faltered^[9, 10]^. Chimeric Antigen Receptor (CAR)-T cell therapy stands out as a pioneering success in this domain, particularly in the treatment of hematological malignancies^[11, 12]^. CAR-T cells are engineered to express synthetic receptors that can selectively target tumor-associated antigens, thereby inducing a potent and specific anti-cancer immune response^[13]^. Nevertheless, extending the use of CAR-T therapy to solid tumors, including prostate cancer, has been met with significant challenges^[14, 15]^. The complex stromal architecture and immunosuppressive elements within the microenvironment of solid tumors, such as tumor-hijacked immune cells (*e.g*., tumor-associated macrophages) and anti-inflammatory cytokines and chemokines, act in unison to undermine the effectiveness of CAR-T cells, necessitating the exploration of alternative CAR-based approaches capable of penetrating these barriers and improving therapeutic outcomes^[16-18]^.

CAR-engineered macrophages (CAR-Ms) are emerging as a promising alternative to CAR-T cell therapy against solid tumors by harnessing the natural tumor-homing abilities of macrophages^[19-21]^. Macrophages are inherently proficient at navigating the complex tumor stroma and infiltrating the tumor microenvironment^[22]^. The plasticity of macrophages allows for the engineering of CARs that can effectively recognize and phagocytose cancer cells^[19, 23]^. Additionally, CAR-Ms have the capacity to emit pro-inflammatory cytokines, which can recruit and activate other immune cells in the tumor microenvironment, creating a more hostile atmosphere for the cancer^[15, 24]^. Moreover, CAR-Ms act as antigen-presenting cells, displaying neo-antigens to activate T cells, which further amplifies the immune response against the tumors^[25]^. The potential of CAR-Ms lies in their dual ability to directly attack tumor cells and reshape the tumor microenvironment to favor immune-mediated clearance^[26]^.

Prostate-specific membrane antigen (PSMA) serves as an ideal target for CAR-engineered therapies due to its overexpression in prostate cancer cells and limited expression in normal tissues^[27]^. Our strategy focuses on constructing PSMA-specific CAR-Ms that can selectively target and eradicate PSMA-expressing prostate cancer cells^[28, 29]^. By leveraging the targeting capabilities of CAR technology and the phagocytic nature of macrophages, we aim to develop a therapeutic approach that can overcome the limitations of current treatments^[21, 30, 31]^. By assessing the phagocytic activity, cytotoxicity, and cytokine production, our data suggest that PSMA-specific CAR-Ms not only demonstrate robust anti-tumor effects *in vitro* but also show significant therapeutic efficacy *in vivo*, providing a compelling case for further exploration as a novel treatment modality for prostate cancer.

## Results

### Construction of PSMA-specific CAR-Ms

Macrophages are engineered using an anti-PSMA CAR structure mainly encoding anti-PSMA single-chain fragment variable (scFv) and CD3ζ intracellular domain^[21]^ **(Fig. 1A)**. Targeting of prostate cancer is provided by extracellular scFv modules that recognize tumor antigen PSMA. Transmembrane and intracellular modules allow activation of ITAM-mediated downstream signaling. Upon the hypothetical binding of CAR-Ms to antigen-expressing prostate cancer cells, it is anticipated that phagocytosis could be triggered, potentially inducing a proinflammatory program. This could lead to the release of interferon and the upregulation of antigen presentation molecules by CAR-Ms^[21]^ (**Fig. 1B**).

**Fig. 1.**
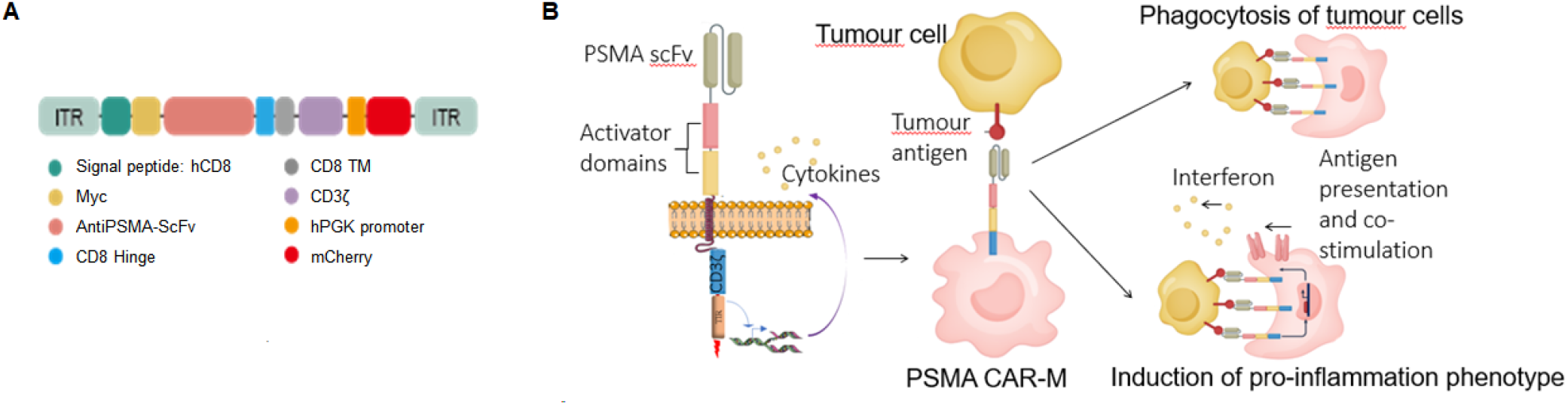
Schematic diagram of PSMA-targeted CAR-M therapy for prostate cancer. **(A)** Structure of PSMA-specific CAR. **(B)** PSMA CAR plasmid was transduced into macrophages by adenovirus. Engineered macrophages stably target PSMA.

### PSMA-specific BMDM CAR-Ms exhibit anti-tumor effects *in vitro*

After transduction of adenovirus carrying PAMS-specific CAR into bone marrow-derived macrophages (BMDM-CAR-PSMA), we confirmed the Myc expression by western blotting (**Fig. 2A**). Flow cytometric analysis showed the BMDM-CAR-PSMA reached 80.7% ratio for positive expression of CAR when compared to BMDMs transduced with adenoviral vector lacking CAR structure (BMDM-Ad) (**Fig. 2B and 2C**). We validated the targeted phagocytosis of CAR-Ms against PSMA-positive human prostate cancer cell line LNCaP *in vitro*. After co-culturing BMDM-CAR-PSMA with LNCaP cells for 6 h, higher phagocytotic efficiency for LNCaP cells was observed for BMDM-CAR-PSMA than that for untreated BMDMs (BMDM-UTD) or BMDM-Ad, respectively (**Fig. 2D**). We then evaluated the killing ability of BMDM-CAR-PSMA, BMDM-UTD or BMDM-Ad against LNCaP cells. We set the effector: target (E: T) ratio to 3:1 for the Fluc-based killing assay. The results showed that the BMDM-CAR-PSMA exhibited a higher killing effect for LNCaP cells, compared to the BMDM-UTD or BMDM-Ad, respectively (**Fig. 2E**). We assessed whether CAR-transduced macrophages exhibited a pro-inflammatory phenotype. The results indicated that pro-inflammatory genes were upregulated, while anti-inflammatory genes did not show significant increases, suggesting that CAR-transduced macrophages were polarized towards a pro-inflammatory state **(Supplementary Fig. 1A and 1B)**. Further analysis during co-cultivation with LNCaP cells revealed that BMDM-CAR-PSMA exhibited a significant upregulation of pro-inflammatory genes compared to BMDM-UTD or BMDM-Ad. In contrast, anti-inflammatory genes did not show significant upregulation. This indicates that BMDM-CAR-PSMA maintains a pro-inflammatory phenotype even within the tumor microenvironment **(Fig.2F and 2G)**. Overall, these findings suggest that PSMA CAR-transduced macrophages are predisposed to a pro-inflammatory state, which could have implications for their efficacy in immunotherapy. Consequently, BMDM-CAR-PSMA were reprogrammed into a pro-inflammatory phenotype and maintained this state in the presence of PSMA-positive prostate cancer cells.

**Fig. 2.**
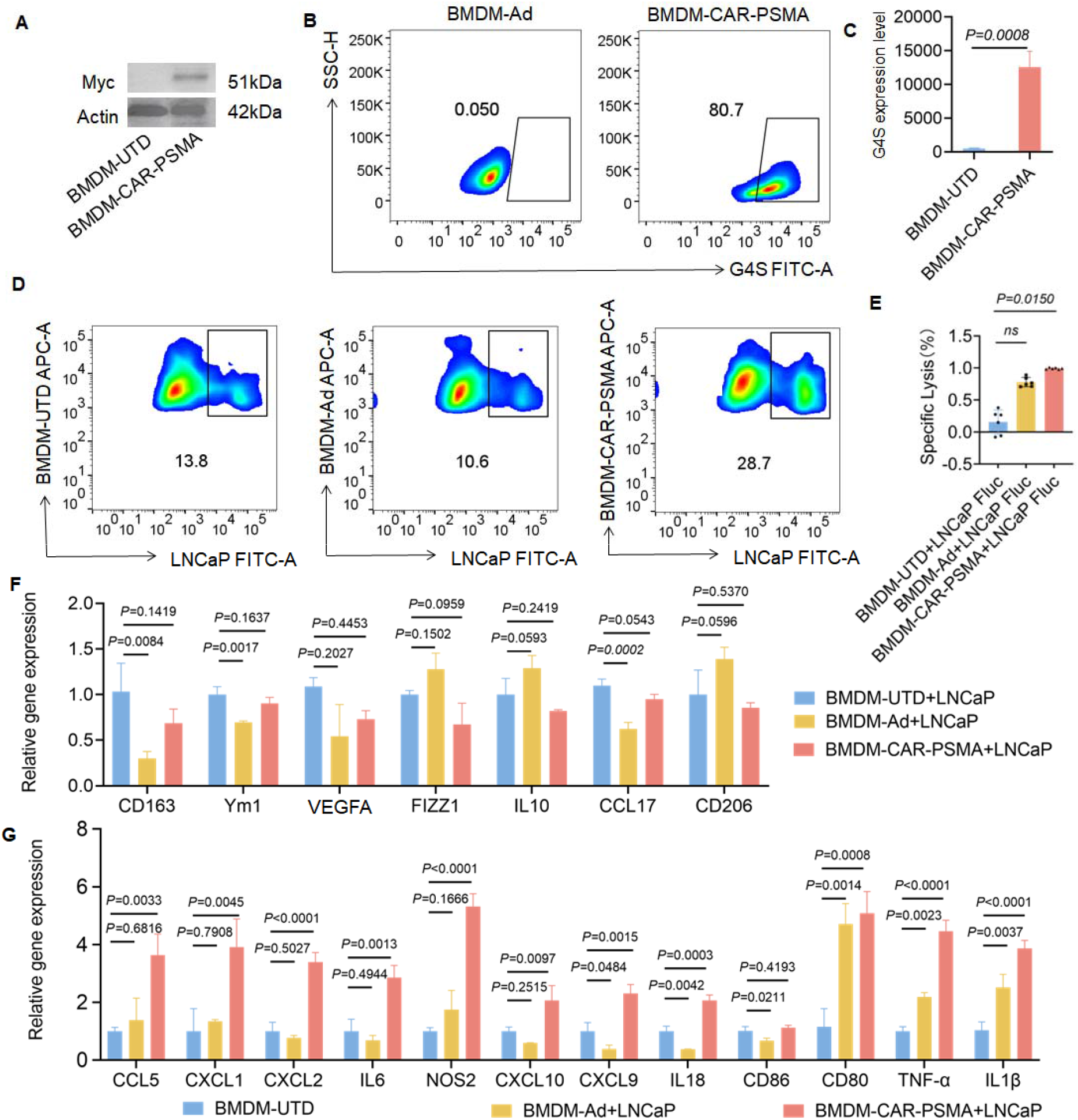
Pro-inflammation phenotype and antitumor effects of BMDM-CAR-PSMA *in vitro*. **(A)** Western blot analysis of Myc-tag protein expression in BMDM-CAR-PSMA and BMDM-UTD cells. **(B)** Flow cytometry for detecting the expression of CAR in BMDM-CAR-PSMA and BMDM-UTD cells. G4S-FITC Linker monoclonal antibody targeted scFv-CAR, and G4S antibody labeled UTD was used as a negative control. **(C)** FACS for determining MFI of CAR expression. Data represent mean ± SD from n=3; statistical significance was calculated with a two-tailed *t-*test. **(D)** Phagocytic effect of BMDM-CAR-PSMA, BMDM-UTD, or BMDM-Ad against LNCaP cells after co-cultivation for 6 h at an E: T ratio of 3:1. **(E)** Killing effect of BMDM-CAR-PSMA, BMDM-UTD, or BMDM-Ad against LNCaP cells after co-cultivation at an E: T ratio of 3: 1 for 24 h. Data are represented as the mean ± SD from n = 5 technical replicates. Statistical significance was calculated using ANOVA multiple-comparison. **(F)** qPCR for detecting the mRNA expression of anti-inflammatory phenotype cytokines after co-culturing BMDM-CAR-PSMA, BMDM-UTD, or BMDM-Ad with LNCaP cells for 6 h with an effector-to-target (E: T) ratio of 1: 3. **(G)** The mRNA expression of pro-inflammation phenotype cytokines was measured after co-culturing BMDM-CAR-PSMA, BMDM-UTD, or BMDM-Ad with LNCaP cells for 6 h with an E: T ratio of 1: 3. Data are presented as mean ± SD from n = 3 technical replicates. Statistical significance was determined using ANOVA multiple-comparison. P < 0.05 was considered statistically significant.

### PSMA-specific CAR-Ms effectively attenuated the progression of tumors *in vivo*

To determine the cytotoxicity of PSMA CAR-M cells *in vivo*, we used a BALB/c nude mouse model bearing xenografts of LNCaP cells (**Fig. 3A**). Sixteen days after LNCaP inoculation, the mice were intravenously injected with BMDM-CAR-PSMA, BMDM-UTD, or BMDM-Ad. The BMDM-CAR-PSMA group showed significantly smaller tumor volume and size than the other groups (**Fig. 3B and 3C**). Hematoxylin-eosin (H&E) staining showed that BMDM-CAR-PSMA induced apoptosis of tumor cells, whereas they did not induce damage to other major tissues including heart, liver, spleen, lung, and kidney (**Fig. 3D and Supplementary Fig. 2A**). Furthermore, immunostaining in tumor tissues indicated that the group treated with BMDM-CAR-PSMA exhibited significantly lower expression of Ki-67, PSMA and TGF-β than the other two groups. Moreover, the BMDM-CAR-PSMA exhibited the highest expression levels of CD68, and IL-1β and enhanced apoptosis among the three groups (**Fig. 3E**). Furthermore, treatment of the mice in BMDM-CAR-PSMA group using PSMA recombinant protein attenuated the antitumor effects of BMDM-CAR-PSMA, indicating the PSMA antigen dependent specificity (**Fig. 3B-3D**). The body weight of the mice showed no obvious alternation among the different treatment groups (**Supplementary Fig. 2B**). Blood biochemical assays demonstrated no acute liver or kidney injury in the mice treated with the CAR-PSMA (**Supplementary Fig. 2C-H**). Collectively, the results of the *in vivo* experiments were consistent with those of the *in vitro*, indicating that BMDM-CAR-PSMA exerted significant anti-tumor effects against prostate cancer in a PSMA-dependent manner.

**Fig. 3.**
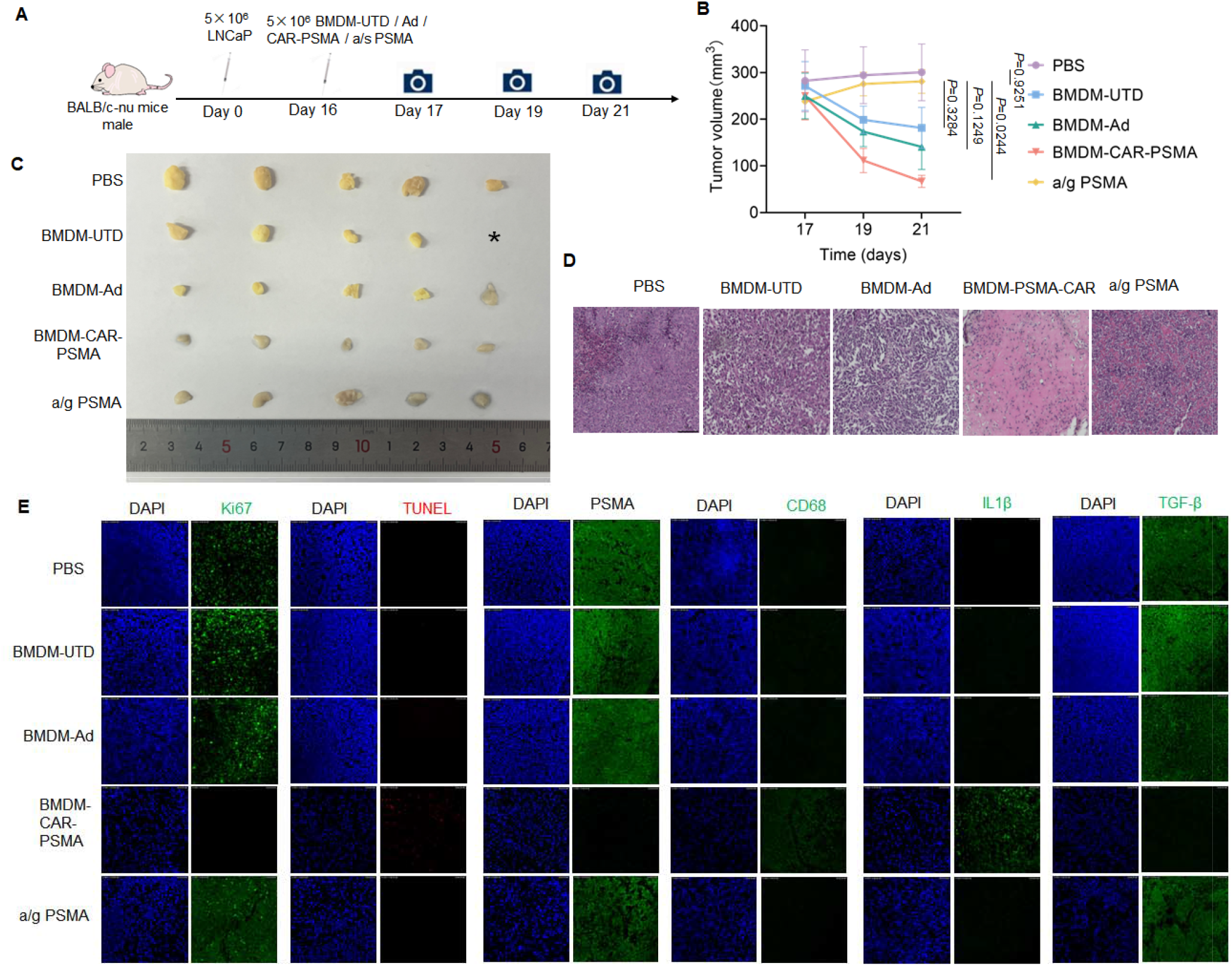
PSMA-specific CAR-Ms effectively attenuated the progression of tumors *in vivo*. **(A)** Illustration of the experimental design for evaluating *in vivo* antitumor efficacy of PSMA-specific CAR-Ms. Briefly, nude mice were subcutaneously injected with LNCaP cells. After 16 days, the mice bearing xenograft tumors were intravenously administrated with PBS, MDM-UTD, BMDM-Ad, BMDM-CAR-PSMA, or BMDM-CAR-PSMA combined with PSMA recombinant protein (a/g PSMA) for 5 days. **(B)** Tumor volumes in each treatment group. Data are presented as mean ± SD (n =5 per group). Statistical significance was determined using one-way ANOVA. **(C)** Macroscopic views of the xenografted tumors from the nude mice in each treatment group. **(D)** Hematoxylin and eosin (H&E) staining of the tumor sections in each treatment group. Scale bar = 50 μm. **(E)** Immunofluorescence staining of the tumor sections for detecting Ki67, CD68, IL-1β, and TGF-β levels, and TUNEL staining for detecting apoptosis in each treatment group. Nuclei were stained with DAPI. Scale bar = 50 μm.

### The hMDM-CAR-PSMA showed anti-tumor ability *in vitro*

Next, adenovirus for PSMA-specific CAR was transduced into human monocyte-derived macrophages (hMDM-CAR-PSMA), and the CAR expression was confirmed by flow cytometry (**Fig. 4A**). After co-culturing hMDM-CAR-PSMA with LNCaP cells for 6 h, we observed that the phagocytotic efficiency of hMDM-CAR-PSMA was significantly higher compared to hMDM-Ad or hMDM-UTD against LNCaP cells (**Fig. 4B**). We conducted an investigation to determine whether CAR-transduced macrophages elicit a pro-inflammatory phenotype. The results revealed that genes associated with pro-inflammatory responses were significantly upregulated, whereas those related to anti-inflammatory processes did not exhibit notable increases, indicating that CAR-transduced macrophages are polarized towards a pro-inflammatory state **(Supplementary Fig. 3A and 3B)**. These findings underscore the presence of a distinct pro-inflammatory phenotype in CAR-transduced macrophages. Moreover, during co-culture experiments, hMDM-CAR-PSMA demonstrated substantial upregulation of pro-inflammatory factors when co-cultured with LNCaP cells, while anti-inflammatory factors showed no significant increase compared to hMDM-Ad or hMDM-UTD **(Fig. 4C and 4D)**. This suggests that hMDM-CAR-PSMA sustains a pro-inflammatory profile within the tumor microenvironment, potentially enhancing its efficacy in targeting cancer cells. Subsequently, we set the effect-target ratio (E: T) to 3:1 for the Fluc-based killing assay, and also confirmed the killing ability of hMDM-CAR-PSMA on LNCaP cells (**Fig. 4E**). Therefore, consistent with the results from BMDM-CAR-PSMA, hMDM-CAR-PSMA exhibited transcriptomic reprogramming towards a pro-inflammatory phenotype and showed a strong killing efficacy against LNCaP cells.

**Fig. 4.**
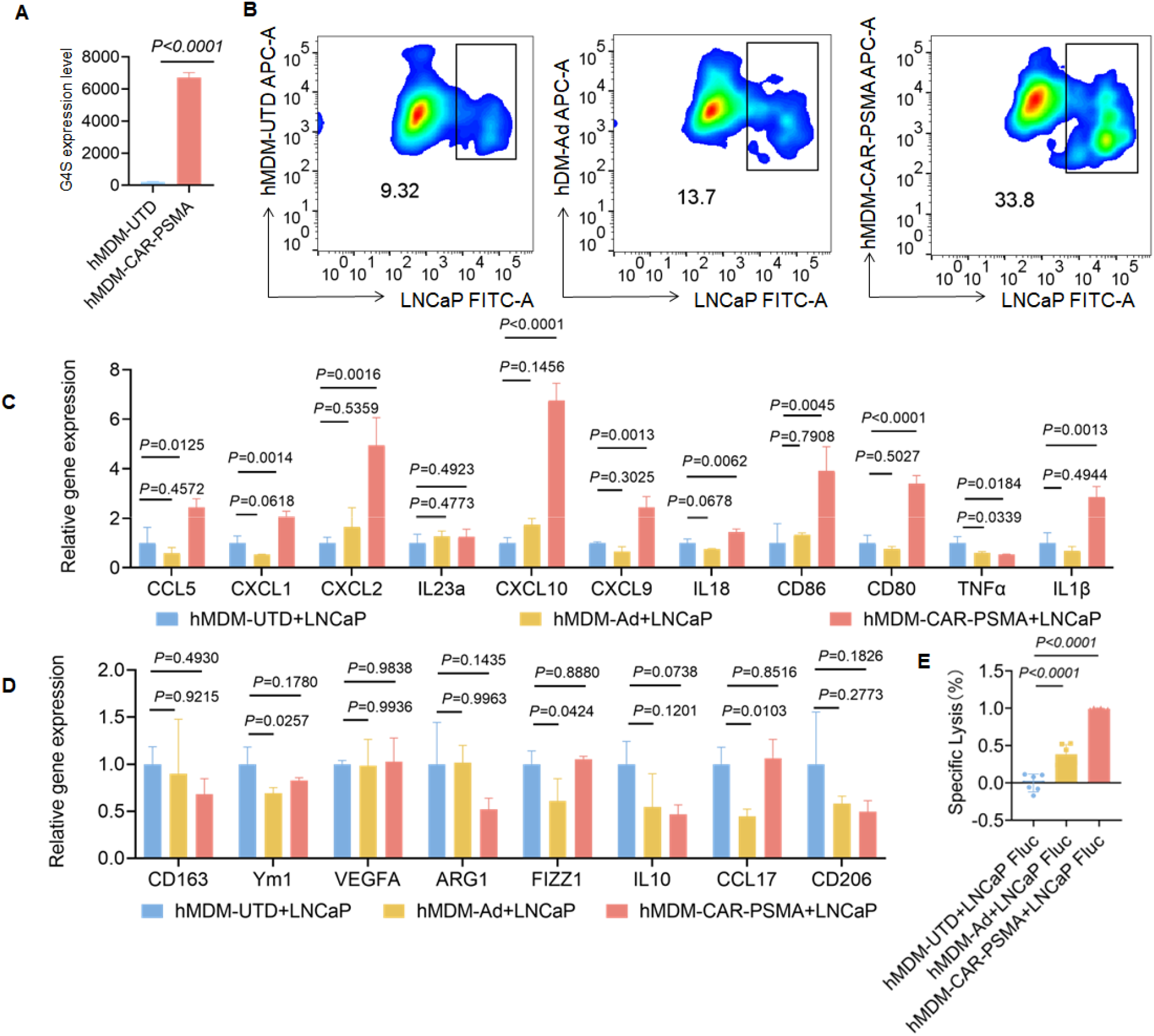
*In vitro* phenotype and antitumor effects of HMDMM-CAR-PSMA. **(A)** FACS for determining MFI of CAR expression. G4S-FITC Linker monoclonal antibody targeted scFv-CAR, and G4S antibody labeled UTD was used as a negative control. Data are represented as mean ± SD of n□= 3. Statistical significance was calculated with a two-tailed *t-*test. **(B)** Phagocytic effect of hMDM-UTD, hMDM-Ad, or hMDM-CAR-PSMA against LNCaP cells after co-cultivation for 6 h at an E: T ratio of 3:1. **(C)** The mRNA expression of pro-inflammation phenotype cytokines after co-culturing of hMDM-CAR-PSMA, hMDM-UTD, or hMDM-Ad with LNCaP cells for 6 h with E: T ratio of 1: 3. **(D)** qPCR for detecting the mRNA expression of anti-inflammatory phenotype cytokines after co-culturing of hMDM-CAR-PSMA, hMDM-UTD, or hMDM-Ad with LNCaP cells for 6 h with E: T ratio of 1: 3. Data are represented as mean ± SD of n□=□3 per treatment group; statistical significance was calculated with a one-way ANOVA. **(E)** Killing effect of hMDM-CAR-PSMA, hMDM-UTD, or hMDM-Ad against LNCaP cells after co-cultivation at an E: T ratio of 3: 1 for 24 h. Data are represented as mean ± SD of n□=□5 per treatment group; statistical significance was calculated with a one-way ANOVA. P < 0.05 was considered statistically significant.

### RNA-seq analysis demonstrated the pro-inflammatory phenotype of hMDM-CAR-PSMA

We conducted RNA-seq analysis for hMDM-CAR-PSMA and hMDM-Ad, respectively. Compared to the hMDM-Ad, a total of 1,565 differentially expressed genes were identified in hMDM-CAR-PSMA, with 859 up-regulated and 706 down-regulated, as shown in the volcano plot **(Fig. 5A)**. Pro-inflammatory genes such as CD80, CXCL2, IL1B, IL23A, CXCL1, CXCL10, IL6, and CD86 were highly expressed, while the anti-inflammatory genes CD163 was significantly down-regulated in hMDM-CAR-PSMA **(Fig. 5B)**. The STRING database was used to construct a protein-protein interaction (PPI) network of differentially expressed genes. Core gene modules were identified, including CXCL10, ISG15, IFIT1, IFIT2, IFIT3, MX1, OAS1, OAS2, and RSAD2 **(Fig. 5C)**. GO enrichment analysis of up-regulated genes in hMDM-CAR-PSMA suggested the activation of pro-inflammatory pathways, such as cytokine activity, receptor-ligand activity, cytokine receptor binding, and chemokine activity **(Fig. 5D)**. However, the down-regulated genes in hMDM-CAR-PSMA were primarily involved in phospholipid binding, aldehyde dehydrogenase (NAD+) activity, aldehyde dehydrogenase [NAD(P)+] activity, and MHC class II protein complex binding (**Supplementary Fig. 4A)**. KEGG analysis demonstrated that the up-regulated genes in hMDM-CAR-PSMA were related to cytokine-cytokine receptor interaction, TNF signaling pathway, IL-17 signaling pathway, rheumatoid arthritis, and JAK-STAT signaling pathway **(Fig. 5E)**. Nevertheless, the down-regulated genes in hMDM-CAR-PSMA predominantly focused on the biosynthesis of unsaturated fatty acids, pyruvate metabolism, arginine and proline metabolism, as well as fatty acid elongation, thereby indicating a significant metabolic shift **(Supplementary Fig. 4B)**. GSEA enrichment analysis showed that pathways such as oxidative phosphorylation and fatty acid metabolism were suppressed, while pathways like inflammatory response were activated in hMDM-CAR-PSMA **(Fig. 5F)**. These findings suggest the upregulation of inflammatory pathways and suppression of metabolic pathways in hMDM-CAR-PSMA, might be the main molecular cascade leading to the therapeutic efficacy against prostate cancer.

**Fig. 5.**
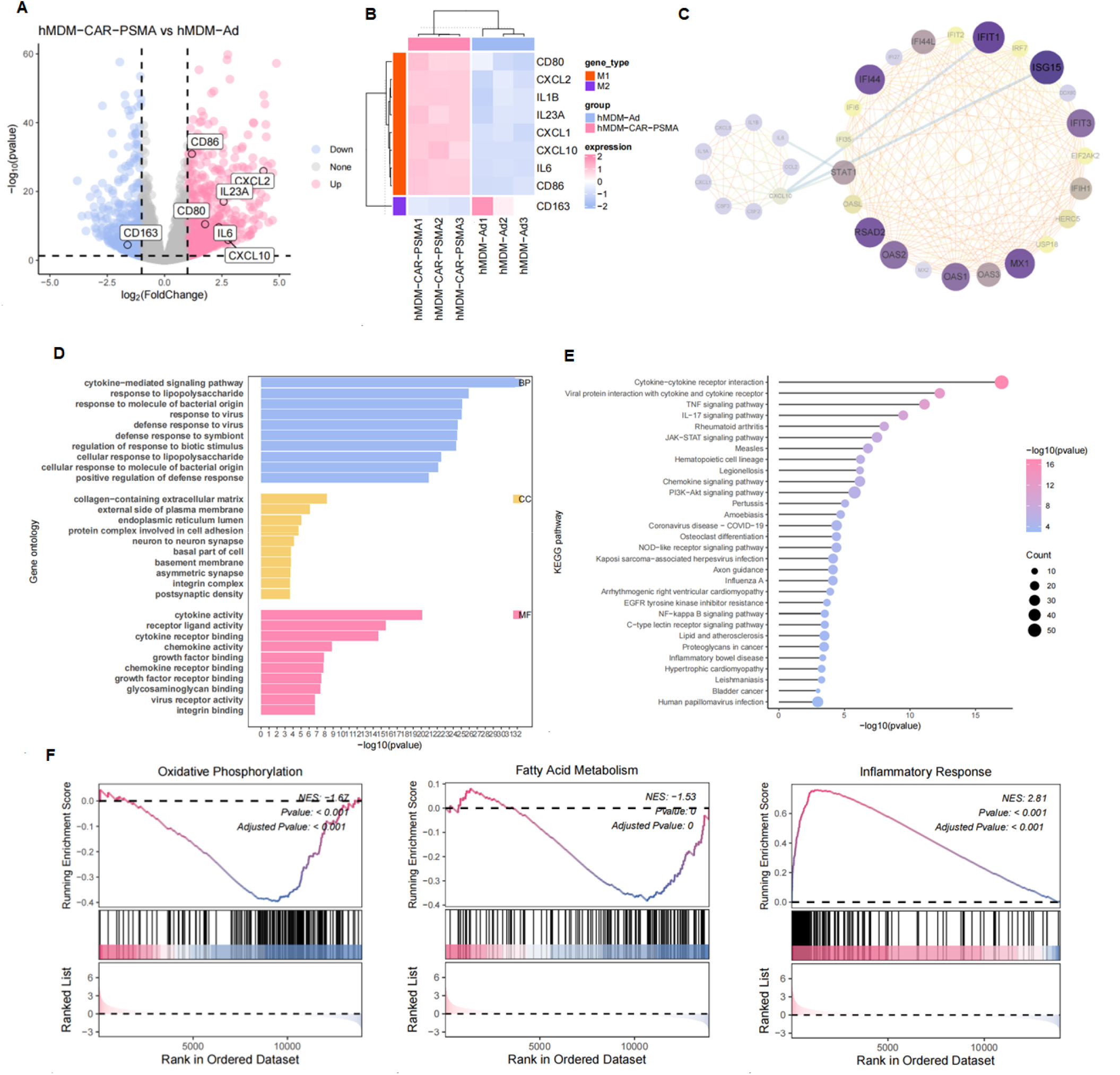
RNA-seq analysis demonstrated the pro-inflammatory phenotype of hMDM-CAR-PSMA. **(A)** Volcano plot showing the distribution of differentially expressed genes, with significant genes detected by qPCR highlighted. The x-axis represents log2 fold change, and the y-axis represents -log10(p-value). Red dots indicate the upregulated genes, blue dots indicate the downregulated genes, and gray dots indicate genes with no significant difference. **(B)** Heatmap displaying differentially expressed macrophage-related genes between hMDM-CAR-PSMA and hMDM-Ad. **(C)** PPI network visualized using Cytoscape software. **(D)** GO pathway enrichment analysis for upregulated genes in hMDM-CAR-PSMA. **(E)** KEGG functional enrichment analysis for the upregulated genes in hMDM-CAR-PSMA. **(F)** GSEA enrichment analysis showing HALLMARK pathway activity.

### Oxidative stress and metabolic changes in hMDM-CAR-PSMA

The RNA-seq analysis indicated a metabolic switch in hMDM-CAR-PSMA cells, and our functional analysis corroborated significant alterations associated with oxidative stress. Notably, the mean fluorescence intensity (MFI) of DCFH was markedly upregulated, signifying elevated intracellular levels of reactive oxygen species (ROS). This increase suggests an influence on mitochondrial function and other metabolic pathways, potentially enhancing the cellular antioxidant capacity to mitigate oxidative damage^[32]^ **(Fig. 6A and 6B)**. The heightened MFI detected via DCFH as a fluorescent probe for ROS reflects increased oxidative stress within these cells, which may be intricately linked to their augmented anti-tumor effects or a more vigorous inflammatory response^[32, 33]^. Further examination of gene expression related to oxidation and glycolysis in hMDM-CAR-PSMA cells was represented in a heat map, illustrating modifications in metabolic pathways that could contribute to the observed enhancement in anti-tumor activity or inflammatory response **(Fig. 6C and 6D)**. Elevated levels of reactive oxygen species (ROS) can instigate a reconfiguration of cellular metabolism by activating inflammatory signaling pathways, resulting in the upregulation of pro-inflammatory cytokines such as IL-1β and CXCL1 **(Fig. 6E)**. This inflammatory response frequently coincides with the downregulation of critical metabolic regulatory genes, including GPR34 and GMPR2, which are indispensable for lipid metabolism and nucleotide biosynthesis, respectively **(Fig. 6F)**. Consequently, this transition towards an inflammatory state may disrupt normal metabolic processes while potentially augmenting the cellular immune response at the expense of metabolic efficiency.

**Fig. 6.**
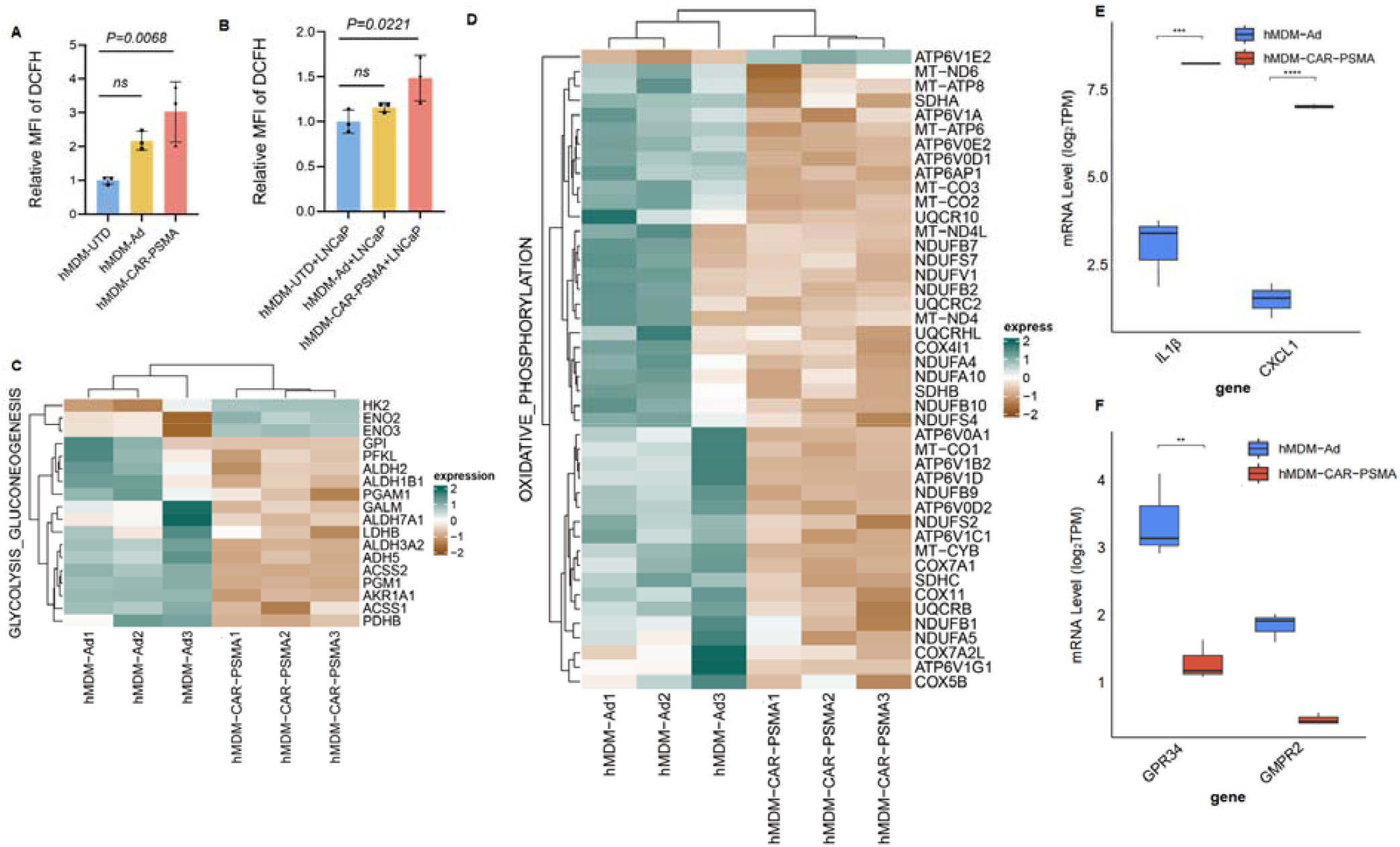
Oxidative Stress and Metabolic Changes Enhance hMDM-CAR-PSMA Efficacy. **(A)** Flow cytometric quantification of relative MFI of 2’,7’-Dichlorodihydrofluorescein (DCFH, n=3) of hMDM-CAR-PSMA. **(B)** Flow cytometry quantification of the relative MFI of 2’,7’-dichlorodihydrofluorescein (DCFH, n=3) was performed after 6 h of co-incubation of hMDM-CAR-PSMA with LNCaP cells. **(C and D)** Heatmap of dysregulated (adjusted p-value <0.05) glycolysis-related **(C)** and amino acid metabolism-related **(D)** gene expression detected in RNAseq of FACS-sorted hMDM-CAR-PSMA after 6 h co-incubation with LNCaP cells. **(E and F)** Two significantly up-regulated genes**(E)** and two down-regulated genes**(F)** were selected from RNAseq data and analyzed by line box plots.

## Discussion

Recent advancements in immunotherapy have highlighted the potential of CAR to reprogram macrophages, overcoming their natural resistance to genetic modification and guiding them toward a pro-inflammatory phenotype^[34]^. This study focuses on leveraging this capability by introducing a PSMA-specific CAR-M and evaluating its therapeutic impact on prostate cancer. CAR-Ms are engineered to infiltrate the tumor microenvironment, target specific tumor antigens, directly phagocytize cancer cells, remodel the tumor microenvironment, and activate other immune cells through enhanced antigen presentation^[10, 35]^. By reprogramming macrophages to the pro-inflammation phenotype, CAR-M therapy seeks to counteract the immunosuppressive environment commonly associated with advanced prostate cancer, enhancing the immune response and promoting tumor destruction^[26]^.

Prostate cancer has been described as an immunological desert with few T cell recruitment. Histopathological analysis confirmed the enriched infiltration of PSMA CAR-M in tumors with reduced tumor cell density, decreased expression of the proliferation marker Ki67, and diminished PSMA expression. Moreover, increased TUNEL staining, CD68 expression, and IL-1β levels in PSMA-specific CAR-M-treated tumors suggest that these engineered macrophages induced tumor cell apoptosis and recruited additional immune effectors, further amplifying the anti-tumor response.

Transcriptional analysis showed that the elevated pro-inflammation gene expression in PSMA CAR-Ms, and persistence of this unique state under co-culture condition with tumor cells, indicating the repressive tumor environment is not able to reprogram the PSMA CAR-Ms. Gene expression profiling revealed that PSMA-specific CAR-M were enriched in pro-inflammatory pathways, with concomitant downregulation metabolic processes associated with the anti-inflammatory phenotype. These metabolic and functional changes are expected to enhance macrophage anti-tumor activity by sustaining inflammation and promoting immune activation within the tumor.

In our analysis of hMDM-CAR-PSMA cells, we observed a significant upregulation of DCFH MFI, indicating increased oxidative stress and ROS levels, which is linked to enhanced anti-tumor effects and a stronger inflammatory response. Metabolic pathway analysis and gene expression heat maps showed changes in oxidation and glycolysis, further supporting these effects. Line box plot analysis of key genes (IL1B, CXCL10, GPR34, GMPR2) confirmed these findings. Collectively, these results underscore the role of oxidative stress and metabolic reprogramming in driving the superior anti-tumor efficacy of hMDM-CAR-PSMA cells.

The clinical translation of PSMA CAR-M therapy may offer a new therapeutic option for patients with prostate cancer, particularly those resistant to conventional treatments. The capacity of PSMA CAR-Ms to home to tumor sites, as demonstrated by their accumulation in inflamed tissues along with off-target effects in non-tumor tissues, and their subsequent polarization towards a pro-inflammation phenotype, provides a dual mechanism of action: direct tumor cell elimination via phagocytosis and indirect suppression of tumor progression through modulation of the immune landscape within the tumor environment.

In conclusion, our study lays the groundwork for the development of PSMA-specific CAR-M as a novel innate immune cell therapy for prostate cancer. The dual functionality of these cells-encompassing both direct tumor cell cytotoxicity and immune modulation-presents a promising strategy for overcoming current therapeutic limitations. Moving forward, continued optimization of CAR-M design, supported by rigorous preclinical and clinical evaluations, will be essential to advance this innovative therapy toward clinical application for prostate cancer treatment.

## Materials and Methods

### Plasmid construction and virus

For replication-deficient adenovirus production, the anti-PSMA CAR was cloned into the pShuttle transfer plasmid using Xba-I and Sal-I, and subsequently cloned into pAd5/F35 using I-Ceu I and PI-Sce I. All cloning steps were validated by restriction enzyme digest and sequencing. The pAd5/F35 adenoviral backbone adenoviral vector lack the E1A, E1B and E3 genes and was made to contain the mCherry signature. Ad5/F35-CAR-PSMA-CD3ζ was generated, expanded, concentrated and purified using standard techniques in HEK-293A cells by the Applied Biology Laboratory, Shenyang University of Chemical Technology. All adenoviral batches were verified negative for replication-competent adenovirus and passed sterility and endotoxin analysis. Adenovirus titers were estimated by direct measurement of the optical density of viral particles using ultra-purified adenovirus (OD_260_) and validated by functional transgene expression in BMDMs and hMDMs. The multiplicity of infection (MOI) was set to 1000 plaque-forming units (PFU) per cell, unless otherwise specified.

### Cell culture

293FT and HEK-293A cells were cultured in Dulbecco’s modified Eagle medium (DMEM, Corning), and RAW264.7 cells in RPMI 1640 (Corning), all sourced from American Type Culture Collection (ATCC). Mouse BMDMs were isolated from C57BL/6 mice femurs and tibias, then differentiated into M0 macrophage with 25 ng/ml recombinant macrophage colony-stimulating factor (M-CSF) (R&D Systems, Minneapolis, MN, USA) for 4 days. The medium contained 10 % FBS (Excel), 100 μg/ml Penicillin-Streptomycin (Sangon), and 100 μg/ml Glutamine (Sangon). Adenovirus was added on day 5 at an MOI of 5×10^3^. Differentiated Macrophages were harvested on day 7 for CAR expression, differentiation, and macrophage purity analysis by Fluorescence Activated Cell Sorting (FACS). Peripheral blood mononuclear cells (PBMCs) were obtained from buffy coats of normal human donor blood provided by the Chinese PLA General Hospital. PBMCs were isolated via gradient density centrifugation. Briefly, 2 ml of blood mixed with 2 ml of PBS was layered over 5 ml of density reagent and centrifuged at 600 g for 20 minutes. The PBMC layer, located just above the density reagent, was carefully collected, washed with 10 ml of PBS, and centrifuged at 700 g for 10 minutes. Cells were counted using a haemocytometer, aliquoted, frozen, and stored in liquid nitrogen. CD14+ monocytes were isolated from PBMC by positive selection using magnetic beads (Miltenyi Biotech) and seeded in RPMI 1640 (Corning) supplemented with 10 % FBS (Excell), 100 μg/ml Penicillin-Streptomycin, and 100 ng/mL M-CSF (PeproTech). Human monocyte-derived macrophages (hMDMs) were seeded in six-well plates with RPMI, 10% FBS, penicillin, streptomycin, 1× GlutaMAX, 1× HEPES and 10ng ml^−1^ recombinant Human M-CSF (300-25, PeproTech) for 7 days. Differentiation was monitored by light microscopy. Adenovirus was added on day 5 at a MOI of 5×10^3^. Differentiated macrophages were harvested on day 7 for CAR expression, differentiation, and macrophage purity analysis by FACS. For smaller-scale experiments, macrophages were plated directly and transduced at a MOI of 5,000 PFU in well plates or flasks. All cells were cultured at 37° C and 5 % CO2.

### Western blot assay

Following various treatment, Cell lysates and viral samples were supplemented with 4x loading buffer and run on denaturing SDS-PAGE. Proteins were then transferred to a PVDF membrane, which was first incubated with a blocking solution, followed by incubation with primary antibodies (1:5000). After washing with PBST, the membrane was incubated with a secondary antibody (Beyotime), washed again, and scanned using the ChemiDoc Touch Imaging System (BIO-RAD). The primary antibodies used were as follows: mouse anti-PSMA (Cell Signaling Technology), mouse anti-V5-Tag (Cell Signaling Technology), rabbit anti-Actin (Beyotime), and mouse anti-myc (Abcam).

### Flow cytometry

Following adenovirus transduction, adherent BMDMs were washed twice with PBS, incubated with Trypsin-EDTA (Gbico) at 37 °C for 5 min, centrifuged at 300 g for 3.5 min, and resuspended in PBS. The cells were then analyzed for red fluorescence emission at 610 nm (APC) using flow cytometry (BD FACSCanto SORP, BD Biosciences). hMDMs were processed and analyzed using the same procedure. LNCaP adherent tumor cells were labled with CellTracker ™ Green CMFDA (ThermoFisher), washed twice with PBS, and incubated with Trypsin-EDTA (Gibco) at 37 °C for 5 min. The cells were then centrifuged at 300 g for 3.5 min, resuspended in PBS, and analyzed for green fluorescence luminescence at 510 nm (FITC) using flow cytometry. Flow cytometry data were further analyzed using FlowJo.

### Fluorescence Activated Cell Sorting

Macrophages derived from 1×10^5^ UTD, control, or CAR-PSMA-CD3ζ mouse monocytes (48 h after transduction) were co-cultured with 3×10^5^ LNCaP (green labeled) target cells for 3 h at 37 ° C in triplicate. After co-culture, BMDMs were washed twice with PBS, incubated with trypsin-EDTA (Gbico) at 37 ° C for 5 min, centrifuged at 300 g for 3.5 min, and resuspended in PBS. BMDMs were then sorted from the target cells using fluorescence-activated Cell Sorting (BD FACSCanto SORP, BD Biosciences) by detecting red fluorescence emission at 610 nm (APC).

### Quantitative real-time PCR

To detect BMDM polarization, BMDMs were inoculated into two sets of six-well plates and infected with either pAd5/F35-CAR-CD3ζ, or pAd5/F35-CAR-PSMA-CD3ζ. One group was infected with adenovirus alone, while the other group was infected with adenovirus and then co-culture with LNCaP cells. After co-culture, BMDMs were sorted out by FACS (FACSAria SORP, BD Biosciences). RNA was isolated using Eastep Super Total RNA Extraction Kit (Promega). cDNA was synthesized using Transcriptor reverse transcriptase (TaKaRa) primed with oligo(dT). The real-time PCR was performed on a CFX96 Real-Time PCR detection system (Bio-Rad) in a 25 μL reaction containing TBGreen™ Premix Ex Taq™ II (Tli RNaseH Plus) (TaKaRa), 0.4 μM of each forward and reverse primer, and RT product diluted at least 10 × (**Supplementary Table 1**). The same method was used to detect hMDMs polarization.

### RNA sequencing

RNA sequencing was performed on RNA isolated from human macrophages obtained from volunteer donors and treated with pAd5/F35-CAR-CD3ζ and pAd5/F35-CAR-PSMA-CD3ζ adenovirus. RNA extraction was carried out using TRIzol. After sequencing, the raw reads were filtered t<u>o</u> remove adaptor sequences, contamination, and low-quality reads. First, raw data containing adapter sequences or low-quality sequences were filtered out. Data processing was then carried out to remove contamination and ensure the integrity of the valid data, which was performed using R Studio. The cleaned reads were subsequently mapped to the human genome using STAR. Gene counts were calculated using featureCounts. Differentially expressed genes were identified by employing the Deseq2 package. We regarded genes with a fold change (FC) greater than 2 or less than -2 as significantly differentially expressed, along with a P-value of 0.05. Additionally, Gene Ontology (GO) and Kyoto Encyclopedia of Genes and Genomes (KEGG) pathway analysis were conducted by R package clusterProfile.

### Animal studies

All mouse studies were conducted in accordance with national guidelines for the humane treatment of animals and were approved by the Institutional Animal Care and Use Committee (IACUC) at the Southern University of Science and Technology. A subcutaneous transplantation tumor model in nude mice was established by injecting LNCaP cells mixed with Matrigel into the right lower limb. 5×10^6^ BMDM macrophages were mixed in 200ul of PBS and administered via intravenous (IV) tail vein injection. Mice were weighed weekly and were subject to routine veterinary assessment for signs of overt illness. Animals were killed at experimental termination or when predetermined IACUC rodent health endpoints were reached.

### Immunofluorescence (IF) assay

To assess the cytotoxicity of BMDM cells and PSMA expressions in subcutaneous tumor models, immunostaining was performed on formalin-fixed, paraffin-embedded tumor tissue sections. The following primary antibodies were used: mouse monoclonal anti-KI67 antibody (1:400, Cell signaling technology), mouse monoclonal anti-CD68 antibody (1:200, MCE), mouse monoclonal anti-IL-1β antibody (10ug /ml, Abcam) and mouse polyclonal PSMA recombinant protein (2 µg/ml Thermofisher). The paraffin-embedded tumor biopsy samples were also subjected to detection using the Colorimetric TUNEL Apoptosis Assay Kit, following biotin labeling and subsequent steps including DAB chromogenic staining. To observe the distribution and morphology of different cell types in tissues, paraffin-embedded sections of tumors, heart, liver, spleen, lung, and kidney samples were stained using the Hematoxylin-Eosin (H&E) Stain Kit (Solarbio). Imaging was performed using an SP8 X confocal microscope (TCS SP8, Leica) equipped with a ×20 lens.

### DCFH reactive oxygen species detection

The hMDM-CAR-PSMA cells were resuspended in PBS and DCFH-DA(2’, 7’ -dichlorodihydrofluorescein diacetate) was added to a final concentration of 10µM. The cells were incubated at 37 ° C for 30 min in the dark to allow dye uptake and ROS detection. After incubation, cells were washed with PBS to remove unincorporated DCFH-DA. The washed cells were re-suspended in PBS and analyzed using flow cytometry. The fluorescence intensity of oxidized DCFH (DCF) was measured to assess ROS levels. The mean fluorescence intensity (MFI) of DCF was recorded. The increase in MFI indicates elevated ROS levels in HMDMM-CAR-PSMA cells. Unstained cells were used as negative controls and cells treated with known ROS inducers were used as positive controls to ensure the accuracy and reliability of the results.

### Serum assays

Following treatment, serum was collected for biochemical analysis. Liver enzymes (ALT, AST, ALP), kidney function markers (creatinine, urea), and serum proteins (albumin, total protein) were measured using an automatic biochemistry analyzer (MS-480, MedicalSystem).

### Statistics

GraphPad Prism software was used for graph generation and statistical analyses. Data are presented as mean ± standard deviation (SD). Comparisons between two groups were conducted using a two-sided unpaired *t-*test. For comparisons among multiple groups, one-way ANOVA followed by Dunnett’s test was employed. Differences in tumor volume and body weight between groups were analyzed using two-way ANOVA followed by Tukey’s multiple comparison test. P < 0.05 was considered statistically significant.

## Acknowledgments

The authors acknowledge the assistance of the Southern University of Science and Technology Core Research Facilities, the Microscope and Imaging Center, and the Experimental Animal Center of the Southern University of Science and Technology. This work is supported by the National Natural Science Foundation Council of China (82472394, 82172386 and 81922081 to C.L., and 32371490 to L.Z.), the Department of Education of Guangdong Province (2021KTSCX104 to C.L.), the Guangdong Basic and Applied Basic Research Foundation (2022A1515012164 to C.L.), the Science, Technology and Innovation Commission of Shenzhen (JCYJ20210324104201005 to C.L.), and supported by Shenyang University of Chemical Technology-Roc Rock Biotechnology (Suzhou) Co., Ltd joint research grant (2021210101004606), and supported by 2023 Ganzhou City Technology Innovation Center [2023ZXGX7956], and 2023 National Center for Biotechnology Innovation in Cellular Therapy “Unveiling and Commanding” Technology Research Project - Macrophage Drug Development Targeting Solid Tumors [2023XB02008] to X.Y.

## AUTHOR CONTRIBUTIONS

X.Y, C.L. and L.Z. conceived the study. X.Y., C.L. L.Z. and Y.X. designed key experiments. Y.X., D.X., Y.Z. and Y.J. carried out experiments. C.C. performed bioinformatics analysis. Y.X. wrote the manuscript. X.Y., C.L. and L.Z. revise the manuscript. X.Y., C.L. and L.Z. provided supervision and funding for the study.

## CONFLICTS OF INTEREST

A patent application has been filed based on partial results presented here for the treatment of prostate cancer with PSMA targeted CAR-M cells [202411225593.1].

