## Supplementary Figure 2 for "PSMA-specific CAR-engineered macrophages for therapy of prostate cancer"

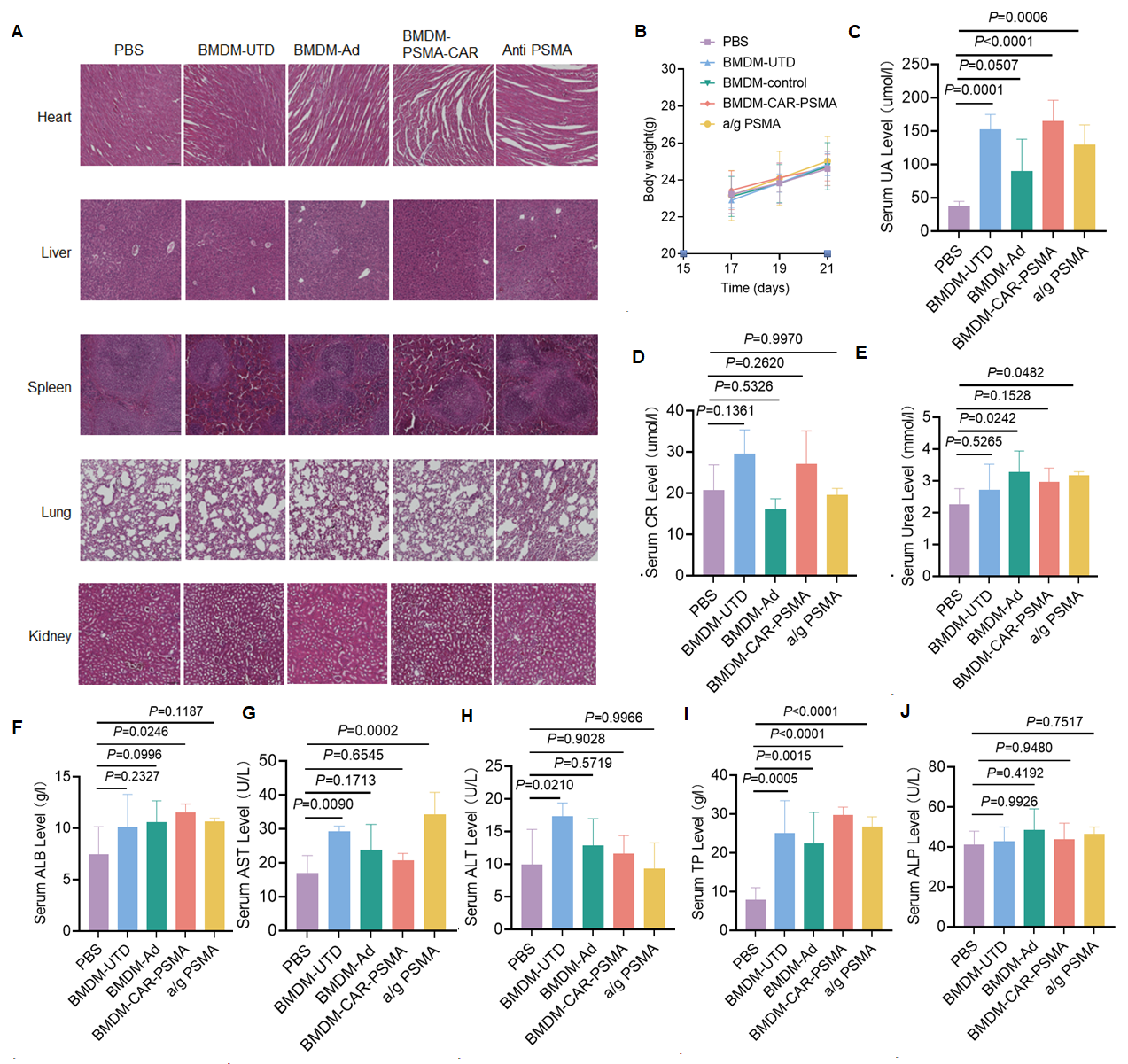

**Supplementary Fig. 2 Safety of BMDM-CAR-PSMA *in vivo*.**

**(A)** Histological assessments of major organs of the mice in each treatment group by H&E staining. Scale bars, 50 μm. **(B)** Body weight of nude mice subcutaneously transplanted with LNCaP cells in each group. **(C-E)** Renal function index including uric acid **(C)**, creatinine **(D),** and Urea **(E)** of the mice in each treatment group. For all panels, data are represented as mean ± SD of n = 5 per treatment group; statistical significance was calculated with one-way ANOVA. ns, no significance. **(F-J)** Liver function index including aspartate Albumin**(F)**, aminotransferase **(G)**, alanine aminotransferase **(H)**, total protein **(I)** and alkaline phosphatase **(J)** of the mice in each treatment group. For all panels, data are represented as mean ± SD of n = 5 per treatment group; statistical significance was calculated with one-way ANOVA. P < 0.05 was considered statistically significant.
