## Supplementary Figure 3 for "PSMA-specific CAR-engineered macrophages for therapy of prostate cancer"

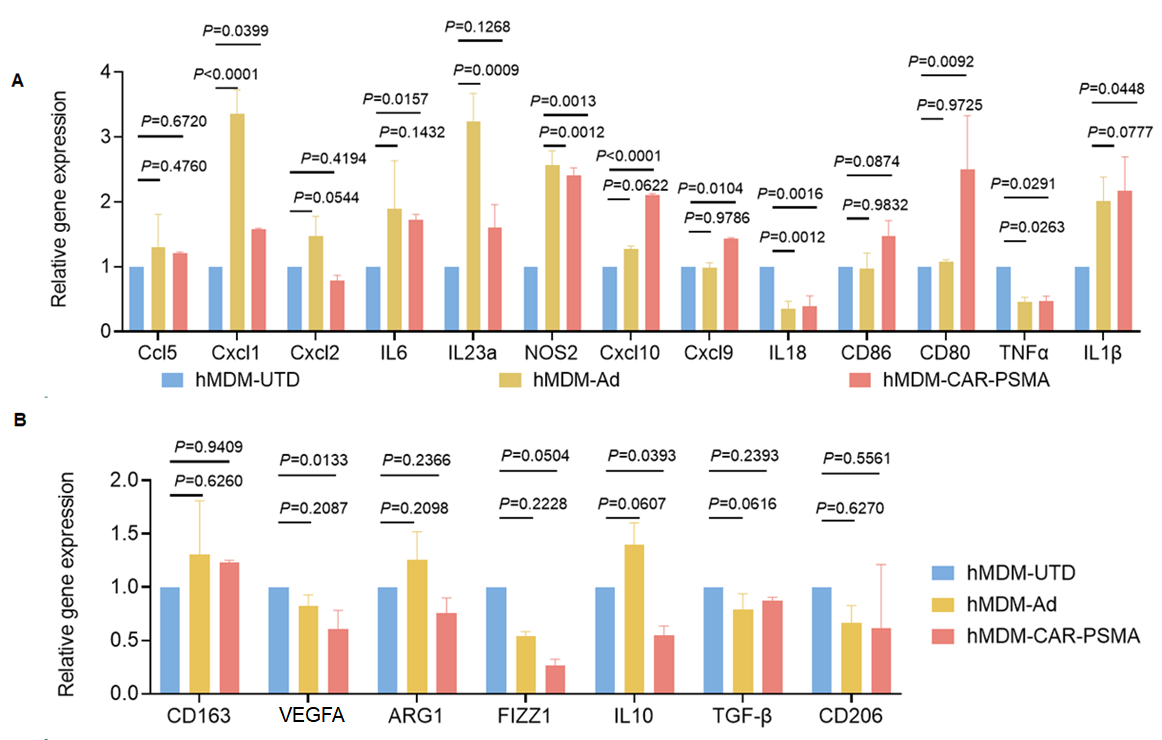

**Supplementary Fig. 3 The mRNA expression of hMDM-CAR-PSMA**

**(A)** qPCR for detecting the mRNA expression of pro-inflammation genes in hMDM-CAR-PSMA, hMDM-UTD, or hMDM-Ad. **(B)** qPCR for detecting the mRNA expression of anti-inflammatory genes in hMDM-CAR-PSMA, hMDM-UTD, or hMDM-Ad. Data are represented as mean ± SD of n = 5 per treatment group. Statistical significance was calculated with a one-way ANOVA. P < 0.05 was considered statistically significant.
