## Supplementary Figure 4 for "PSMA-specific CAR-engineered macrophages for therapy of prostate cancer"

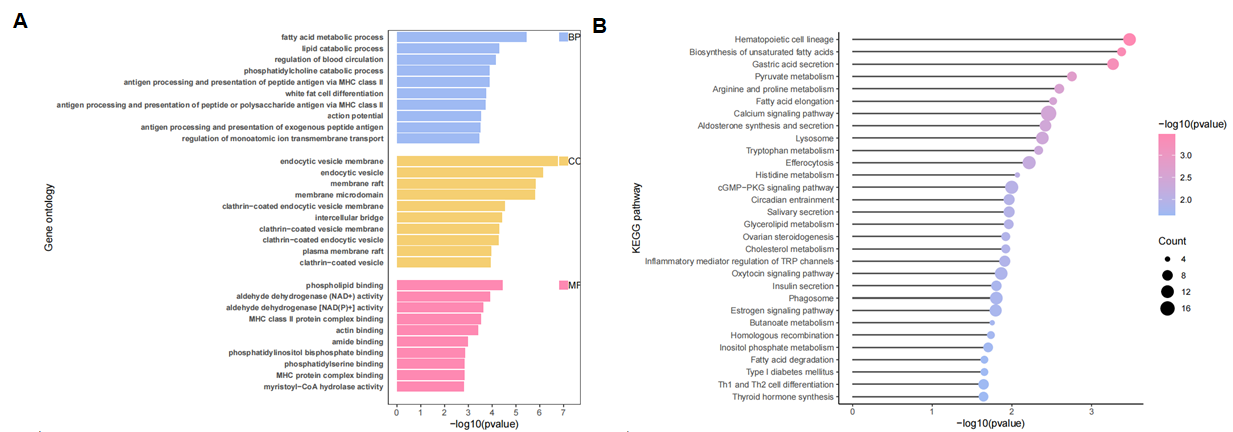
 **Supplementary Fig. 4 GO pathway enrichment and KEGG analysis.**

**(A)** GO pathway enrichment analysis for the downregulated genes in BMDM-CAR-PSMA. **(B)** KEGG functional enrichment analysis for the downregulated genes in BMDM-CAR-PSMA.
